# The “adductome”: a limited repertoire of adducted proteins in human cells

**DOI:** 10.1101/872176

**Authors:** Kostantin Kiianitsa, Nancy Maizels

## Abstract

Proteins form adducts with nucleic acids in a variety of contexts, and these adducts may be cytotoxic if not repaired. Here we apply a proteomic approach to identification of proteins adducted to DNA or RNA in normally proliferating cells. This approach combines RADAR fractionation of proteins covalently bound to nucleic acids with quantitative mass spectrometry (MS). We demonstrate that “RADAR-MS” can quantify induction of TOP1- or TOP2-DNA adducts in cells treated with topotecan or etoposide, respectively, and also identify intermediates in physiological adduct repair. We validate RADAR-MS for discovery of previously unknown adducts by determining the repertoires of adducted proteins in two different normally proliferating human cell lines, CCRF-CEM T cells and GM639 fibroblasts. These repertoires are significantly similar with one another and exhibit robust correlations in their quantitative profiles (Spearman r=0.52). A very similar repertoire is identified by the classical approach of CsCl buoyant density gradient centrifugation. We find that in normally proliferating human cells, the repertoire of adducted proteins — the “adductome” — is comprised of a limited number of proteins belonging to specific functional groups, and that it is greatly enriched for histones, HMG proteins and proteins involved in RNA splicing. Treatment with low concentrations of formaldehyde caused little change in the composition of the repertoire of adducted proteins, suggesting that reactive aldehydes generated by ongoing metabolic processes may contribute to protein adduction in normally proliferating cells. The identification of an endogenous adductome highlights the importance of adduct repair in maintaining genomic structure and the potential for deficiencies in adduct repair to contribute to cancer.

## 1. Introduction

Protein-nucleic acid adducts can form in the course of enzymatic reactions or as a result of treatment with agents that cause proteins to become crosslinked to DNA or RNA. Adducts can be cytotoxic, and robust pathways carry out adduct repair [1-3]. More than 30 human proteins form transient adducts with DNA as obligatory reaction intermediates (**Table S1**), among them topoisomerases, methyltransferases, tyrosyl-DNA phosphodiesterases, DNA glycosylases, polymerases and repair proteins with AP lyase activity [1, 4]. Treatment of cells with chemicals or radiation causes a much wider spectrum of proteins to become crosslinked to nucleic acids, and details of processes essential to DNA replication, repair, transcription, RNA processing, and translation have been elucidated by combining chemical and UV crosslinking with precise characterization of interacting sites and motifs (e.g. [5-7]).

Formaldehyde crosslinking is one of the most common experimental approaches for mapping intracellular molecular interactions. Formaldehyde readily permeates mammalian cell membranes and stimulates crosslinking between a reactive amino group of lysine in close proximity to a nucleic acid substrate. Human cells and tissues generate formaldehyde in the course of normal metabolic processes, such as histone demethylation, repair of alkylated DNA and oxidation of folate [8-10]. The formaldehyde concentration in human plasma exceeds 0.1 mM and may be even higher in some tissues (reviewed by [3]).

The ease with which adducts are induced experimentally by brief treatment with formaldehyde raised the question of whether cells may normally contain some level of adducts formed in response to exposure to endogenous formaldehyde. To address this, we have characterized the repertoire of adducted proteins in normally proliferating human cells, taking advantage of the unbiased detection intrinsic to mass-spectrometry (MS) to identify proteins covalently bound to nucleic acids. Samples were generated for MS by RADAR fractionation [11, 12], a procedure in which adducts are fractionated by cell lysis in chaotropic salts and detergent followed by alcohol precipitation of nucleic acids — both DNA and RNA (**Fig. 1A**). Adducts of human TOP1 and TOP2A, POLβ, E. coli DNA gyrase and other proteins have previously been recovered by RADAR fractionation and quantified by immunodetection [11-18].

**Fig. 1.**
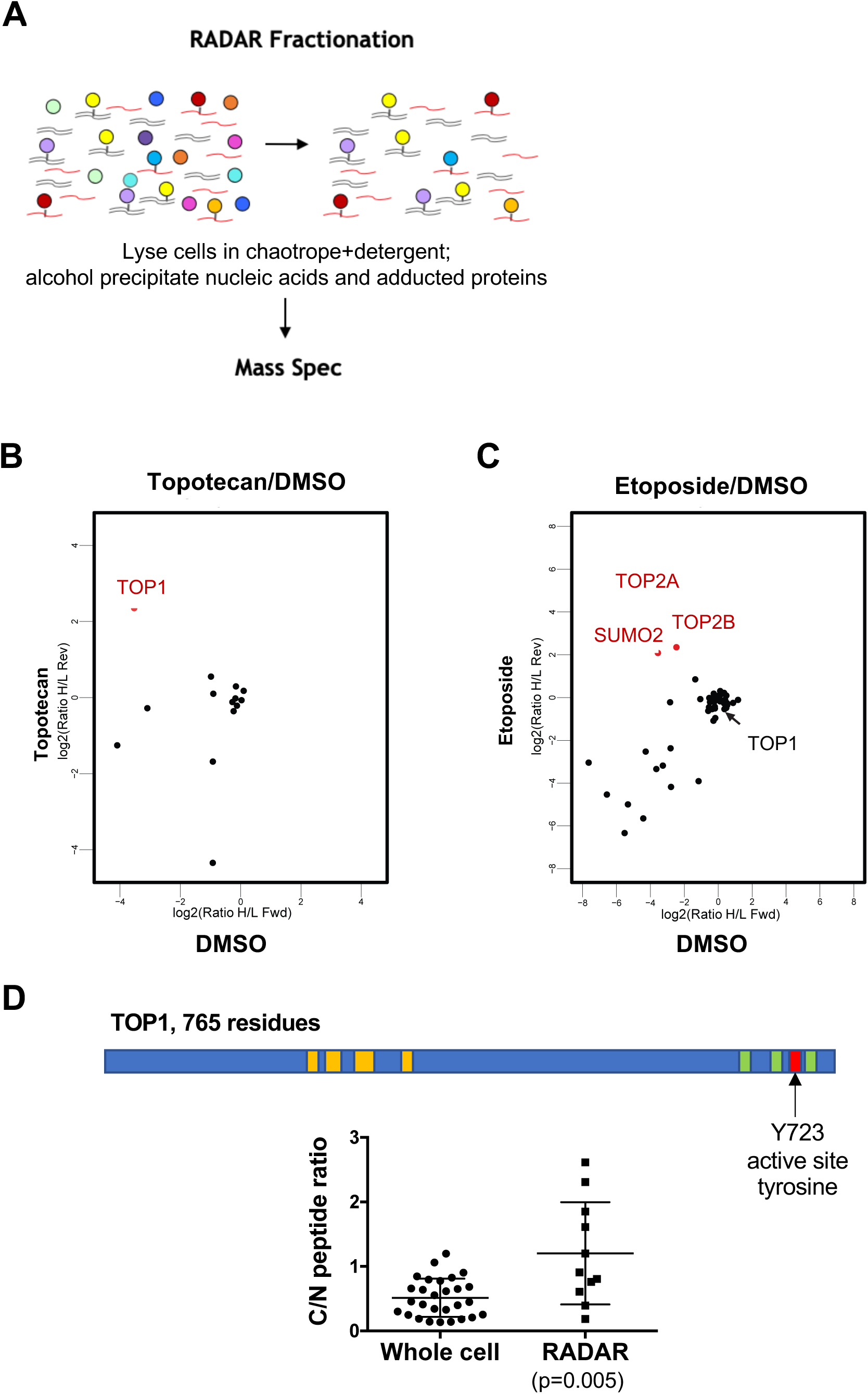
RADAR-SILAC analysis identifies adducts formed by TOP1 and TOP2. (A) Schematic of fractionation: Cells are lysed in chaotropic salts and detergent, then nucleic acids and adducted proteins precipitated with ethanol or isopropanol. (B) RADAR-SILAC analysis of CCRF-CEM cells treated with the TOP1 poison, topotecan. Proteins enriched in inhibitor-treated cells are shown in red. (C) RADAR-SILAC analysis of CCRF-CEM CCRF-CEM cells treated the TOP2 poison, etoposide. Proteins enriched in inhibitor-treated cells are shown in red. (D) *Above*, diagram of TOP1. Tryptic peptides used to quantify recovery from the N-terminal (residues 205-216, 224-239, 252-271 and 300-310,) and C-terminal (residues 643-750, 693-700, 701-712 and 736-742) are highlighted in yellow and green, respectively. Catalytic tyrosine Y723 indicated in red. *Below*, ratios of recovery of the indicated peptides from the C- and N-terminal regions of TOP1, based on intensity. Each dot represents relative recovery in one of 28 different experiments reported in the MaxQB database (whole cell), and squares represent relative recovery in one of 11 different MS analyses of RADAR-fractionated cells. The Mann-Whitney test was used to determine p value.

Here we show that RADAR-MS faithfully identifies TOP1 and TOP2 adducts induced by treatment with topotecan or etoposide, respectively; and also identifies TOP1 fragments that are likely intermediates in physiological repair. We validate RADAR-MS for discovery of previously unknown adducts by determining the repertoires of adducted proteins in two different normally proliferating human cell lines, CCRF-CEM T cells and GM639 fibroblasts. These repertoires are significantly similar with one another and with the repertoire determined using the classical approach of CsCl buoyant density gradient fractionation to recover adducts. We show that treatment with low doses of formaldehyde increases representation of histones and HMG proteins but has little effect on the overall composition of the repertoire of adducted proteins, consistent with the notion that reaction with endogenous formaldehyde contributes to adduction. The repertoire of adducted proteins is enriched for histones, HMG proteins and for proteins involved in RNA splicing. These results suggest that human cells contain an adducted proteome, or “adductome”, comprised of a limited repertoire of proteins in a small number of functional classes. The endogenous adductome may contribute to the burden of adducts requiring repair in order to maintain genomic structure, particularly in cells deficient in adduct repair.

## 2. Materials and methods

### 2.1 Cells, cell culture, drug treatment and SILAC labeling

The CCRF-CEM T lymphoblastoid cell line (ATCC CCL-119), which was derived from a human acute lymphoblastic leukemia, was cultured in RPMI1640 containing 10% fetal bovine serum and Pen-Strep (Gibco) at 37°C in 5% CO_2_. The GM639 cell line, derived from an SV40-transformed human fibroblast, was cultured in DMEM containing 10% fetal bovine serum and Pen-Strep (Gibco) at 37°C in 5% CO_2_. Sensitivity to topoisomerase poisons topotecan and etoposide was demonstrated using the CellTiter-Glo^®^ ATPase assay (Promega). Prior to metabolic labeling analyses, cells were shown to be drug sensitive (**Supplementary Fig. S1**).

For metabolic labeling, light and heavy SILAC RPMI-1640 media were prepared by supplementing RPMI-1640 lacking lysine and arginine with either “light” (normal isotope abundance L-lysine and L-arginine) or “heavy” (stable isotope [^13^C_6_, ^15^N_2_] L-lysine and [^13^C_6_, ^15^N_4_] L-arginine enriched) amino acids [19]. CCRF-CEM cells were metabolically labeled in parallel 40 ml cultures in T-75 flasks at 37°C, 5% CO_2_ in light or heavy SILAC RPMI-1640 media supplemented with 10% dialyzed fetal bovine serum for at least five doublings. When cell density reached 5 × 10^5^ cells/ml, cells were treated for 15 min with 10 *μ*M topotecan (Enzo Life Sciences) or 50 *μ*M etoposide (EMD Biosciences), with controls treated with 0.5% DMSO. Cells (2 × 10^7^) were then harvested by 10 min centrifugation at 2,740 RCF. To validate differences between untreated and treated cultures, label-swap replicates were performed by reversing the SILAC labeled state and the drug treatments between replicates, with the expectation that SILAC ratios for bona fide protein adducts would invert accordingly.

### 2.2 RADAR fractionation

Cells were recovered by centrifugation and lysed in 1 ml of pre-warmed LS1 reagent, consisting of 5 M guanidinium isothiocyanate (GTC), 2% Sarkosyl, 10 mg/ml DTT, 20 mM EDTA, 20 mM Tris-HCl (pH 8.0) and 0.1 M sodium acetate (pH 5.3), adjusted to final pH 6.5 with NaOH. To ensure complete homogenization and reduce viscosity due to high molecular weight DNA, lysates were sonicated on ice using a cuphorn device (QSONICA) or passed 8-12 times through a 22G 1½ inch needle. Each lysate was aliquoted into a pair of Eppendorf tubes, 450 μl/tube (900 μl total), then to each tube was added 150 μl of 8 M LiCl (final concentration 2 M LiCl), followed by an equal volume (600 μl) of isopropanol. Nucleic acids (including bound proteins) were recovered by 10 min centrifugation at 21,000 RCF then washed twice by addition of 1 ml 75% ethanol and 5 min centrifugation. Pellets were dissolved by addition of 100 μl of freshly prepared 8 mM NaOH and through resuspension by shaking on an Eppendorf Thermomixer at room temperature for 15-30 min, then neutralized by addition of 2 μl 1M HEPES per tube. Samples were treated for 30 min at 37°C with RNase A (Fermentas) at final concentration 50 μg/ml. Then 350 μl LS1 reagent and 150 μl 8M LiCl were added, and samples again precipitated with an equal volume (600 μl) of isopropanol. Nucleic acids were recovered and washed twice as above, then resuspended by brief shaking in 50 μl 8 mM NaOH, then neutralized by addition of 1 μl 1M HEPES per tube. DNA and RNA were quantified using DNA/RNA specific fluorescent detection kits (Qubit, Invitrogen). Total protein was measured using a BCA detection kit (Pierce).

### 2.3 Enrichment of adducts by CsCl buoyant density gradient centrifugation

DNA-protein covalent complexes were isolated by ultracentrifugation on a CsCl buoyant density gradient [20]. Briefly, CCRF-CEM cells (2 × 10^7^) were harvested and lysed in 3 ml TE with 1% Sarkosyl supplemented with protease inhibitor cocktail (Roche). The lysate was passed through a 22G 1½ inch needle to reduce viscosity, then loaded on top of a preformed four-step CsCl gradient (1.37, 1.50 1.72, and 1.82 g/ml) and centrifuged in an SW41 rotor at 30,000 rpm for 20 hr at 20°C. Peak fractions containing DNA and DNA-protein adducts were pooled and desalted by diafiltration using a Microcon centrifugal filter (EMD Millipore). This fraction contained 82% dsDNA and 18% RNA, as measured by fluorescent Qubit assays specific for dsDNA and RNA.

### 2.4 NanoLC-MS/MS analysis and quantification

Prior to MS analysis, RADAR fractions in 200 μl volume were treated with 10 units of Cyanase nuclease (RiboSolutions) for 16-18 hr at room temperature, in reactions containing 6 mM MnCl_2_. For SILAC experiments, RADAR fractions from untreated and drug-treated SILAC-labeled cells were mixed in a 1:1 ratio based on DNA concentration before processing. For label-free experiments, each fractionated sample was processed separately. To each sample, 8M urea in 50mM Tris pH 8.0 and 75 mM NaCl was added, followed by reduction and alkylation with Tris2(-carboxyethyl)phosphine (TCEP) and chloroacetamide (CAM) at final concentrations of 1 mM and 2mM, respectively. An equal volume of 100 mM triethylammonium bicarbonate (TEAB; Sigma) was then added and samples were then digested with LysC (Wako Chemicals) at 1:100 ratio for 2 hr. Samples were further diluted to adjust urea concentration below 1.5 M, then digested with Trypsin (Thermo Scientific) at 1:100 ratio for 16-18 hr at room temperature. Peptides were acidified at pH 2.0 with 10% trifluoroacetic acid (TFA) and desalted using StageTips [21]. Peptides were eluted in 80% acetonitrile/0.1%TFA and dried to completion using vacuum centrifugation, then resuspended in 5% acetonitrile/0.1% TFA.

For SILAC experiments with samples from CCRF-CEM and GM639 cells, peptides were separated on a Thermo-Dionex RSLCNano UHPLC instrument (Sunnyvale, CA) with 10 cm long 100 μm I.D. fused silica capillary columns and packed with 3 μm 120 Å reversed phase C18 beads (Dr Maisch), made in house with a laser puller (Sutter). The LC gradient was 90 min of 10-30% B at 300 nL/min. LC solvent A was 0.1% acetic acid and LC solvent B was 0.1% acetic acid, 99.9% acetonitrile. MS data were collected with a Thermo Orbitrap Elite. Data-dependent analysis was applied using Top15 selection with CID fragmentation using 35% collision energy.

For label-free analysis of samples from CCRF-CEM cells, peptides were separated on a Thermo EASY-nLC 1200 UHPLC instrument with in-house packed columns as described above. The LC gradient was 90 minutes of 6-38% B at 300 nL/min. LC solvent A was 0.1% acetic acid and LC solvent B was 0.1% acetic acid, 80% acetonitrile. MS data were collected with a Thermo Orbitrap Fusion Lumos Tribrid. Data-dependent analysis was applied using Top10 selection with simultaneous CID fragmentation at 32% CID collision energy detected in the ion-trap and HCD fragmentation using 31% collision energy and detected in Orbitrap. The resolution of the Orbitrap for is 60,000 for MS1 and 30,000 for MS2.

### 2.5 Data analysis

MaxQuant v.1.5.7.4 and the associated Andromeda search engine was used to search a Uniprot human database (July 2016). The following search parameters were used: Trypsin/P with two missed cleavages, fixed carbamidomethylated cysteines, and variable modifications of oxidized methionines and N-terminal acetylation. Initial FTMS and ITMS MS/MS tolerances were set at 20 ppm and 0.5 Da, respectively. Protein and peptide false discovery rate (FDR) was 1%, minimum peptide length was seven amino acids, and a minimum of two peptide ratios were required to quantify a protein. Data were analyzed with the Perseus and R environments. A published deep whole cell proteome of CCRF-CEM [22] that contained 6,282 unique gene entries with positive IBAQ values was used as a reference for repertoire and peptide intensities of this cell line. To calculate intensities of individual TOP1 peptides recovered from whole cell proteomes, we used a dataset of 11 cell lines deposited in the MaxQB database (maxqb.biochem.mpg.de) [23, 24].

### 2.6 Analysis of gene ontologies

A total of 20,199 entries from the manually annotated and reviewed human proteome (Swiss-Prot) were extracted from the Uniprot (www.uniprot.org). Entries that had no matching gene names (some putative proteins and peptides < 20 amino acids) and non-unique gene names from families with high sequence similarity (e.g. HLA-A) were then eliminated, to generate a list containing 19,742 unique entries. We searched the list of unique entries for gene ontologies related to DNA/RNA/nucleic acids binding properties: GO:0003676 (nucleic acid binding, NBP); GO:0003677 (DNA binding, DBP); GO:0003723 (RNA binding, RBP). This yielded 3,919 NBP, 2,438 DBP and1,577 RBP species. Of the 3919 NBP, 265 were able to bind both DNA and RNA, and 169 lacked a particular DNA/RNA binding GO assignment. As a reference, we used a published deep whole cell proteome of CCRF-CEM that contained 7,625 unique gene entries, of which 6282 had positive iBAQ values [22]. Enriched pathways and gene ontologies were identified using STRING database of protein-protein interaction networks (string-db.org).

### 2.7 Statistical analyses

Hypergeometric distributions were evaluated using the online calculator https://systems.crump.ucla.edu/hypergeometric/index.php. The population sizes, N, and the experimental parameters for each analysis are shown in the corresponding figure legend. For comparisons with whole cell repertoires, the population size was specified at 9000, based on averages of 10,000 proteins per cell and 90% shared repertoire between pairs of human cell lines [23]. Other statistical tests were performed using GraphPad Prism software.

## 3. Results

### 3.1 RADAR-MS detects drug-induced topoisomerase-DNA adducts

TOP1, the most abundant topoisomerase, cleaves and rejoins single DNA strands to regulate superhelicity at the promoter and target repair to non-canonical DNA structures [25]. Topotecan is a derivative of camptothecin, a natural product which stabilizes normally transient TOP1-DNA adducts, thereby generating cytotoxic lesions. CCRF-CEM cells are derived from a human acute lymphoblastic leukemia and are topotecan-sensitive (**Supplementary Fig. S1**). We assayed the ability of RADAR-MS to detect enrichment of TOP1-DNA adducts in CCRF-CEM cells cultured in heavy or light SILAC medium, treated for 15 min with topotecan or DMSO carrier, then RADAR fractionated to recover adducts, nucleic acids eliminated by Cyanase nuclease digestion, and the sample digested with LysC and trypsin to generate peptides suitable for MS analysis. In two independent experiments, TOP1 was consistently found to be enriched in cells treated with topotecan, as shown by a representative plot (**Fig. 1B)**.

TOP2 regulates DNA topology by catalyzing breakage and joining of both DNA strands. Etoposide, a synthetic derivative of a natural toxin, poisons TOP2 by stabilizing normally transient adducts to generate cytotoxic lesions. In two independent experiments, RADAR-MS analysis of cells treated with etoposide documented enrichment of two different human type 2A topoisomerases, TOP2A and TOP2B, as shown by a representative plot (**Fig. 1C)**. As anticipated, TOP1 was not enriched following etoposide treatment. Peptides from SUMO2 were enriched, consistent with previous reports that TOP2A- and TOP2B-DNA adducts are sumoylated, a posttranslational modification that can contribute to proteolytic repair [26, 27]. The SILAC enrichment score of TOP2A was four-fold higher than that of TOP2B, suggesting a higher selectivity of TOP2A poisoning with etoposide in vivo. These results demonstrate that RADAR-MS can identify post-translational modifications and quantitatively discriminate among adducted protein species.

### 3.2 RADAR-MS provides a snapshot of physiological adduct repair

In normally proliferating cells, TOP1 forms transient covalent bonds at some sites, but at AP sites and other endogenous lesions it may form irreversible adducts which must undergo proteolytic repair. This raised the possibility that some of the TOP1 adducts recovered from untreated cells were repair intermediates that had undergone proteolytic processing. The active site tyrosine Y723 in TOP1 that forms stable adducts with DNA is near the C-terminus of the 765 residue TOP1 protein (**Fig. 1D**). If RADAR fractionation recovers partially proteolyzed TOP1-DNA adducts from untreated cells, then C-terminal tryptic peptides of TOP1 will be enriched in RADAR-MS spectra relative to N-terminal peptides. We confirmed this by comparing the relative recovery of N-terminal (residues 205-310) and C-terminal (residues 643-742) tryptic peptides from whole cells in 28 different experiments reported in the MaxQB database with recovery by RADAR-MS analysis as determined in 11 independent experiments. C-terminal peptides were significantly enriched in the RADAR fraction (Mann-Whitney test, p=0.005; **Fig. 1D**).

### 3.3 RADAR-MS identifies a limited repertoire of adducted proteins shared between different cell types

We next sought to characterize species that form endogenous crosslinks with nucleic acids in normally proliferating CCRF-CEM T cells. Two untreated samples (NTa and NTb), each containing 25 × 10^6^ cells, were RADAR-fractionated, yielding 220 and 270 μg DNA, respectively (**Supplementary Fig. S2A**). They were then subject to label-free quantitative MS. The IBAQ intensity of a protein in a MS sample provides a measure of how observed enrichment relates to theoretical enrichment by calculating relative abundance of each protein (its IBAQ score) from the sum of intensities of all identified peptides divided by the number of theoretically observable peptides in its sequence. The MS signal of a sample corresponds to the sum of IBAQ intensities for all identified proteins in that sample. The MS signals for samples NTa and Ntb were 1.9 and 2.4 × 10^10^, respectively (**Supplementary Fig. S2B; Table S2**). Combining the separate repertoires of samples NTa and NTb (1449 and 1511 proteins, respectively) yielded a total repertoire of 1664 distinct proteins, with 1296 shared proteins (**Fig. 2A**). The shared proteins accounted for 78% of the total repertoire and for 99.6% and 98.9% of MS signal in each of the two samples, respectively. Pairwise comparisons of the IBAQ intensities of the 1296 shared proteins showed very high correlation (Spearman r=0.93; p<0.0001; **Fig. 2B**). Thus, RADAR-MS yielded consistent repertoires of adducted proteins, with relative IBAQ intensities of fractionated proteins well reproduced between samples.

**Fig. 2.**
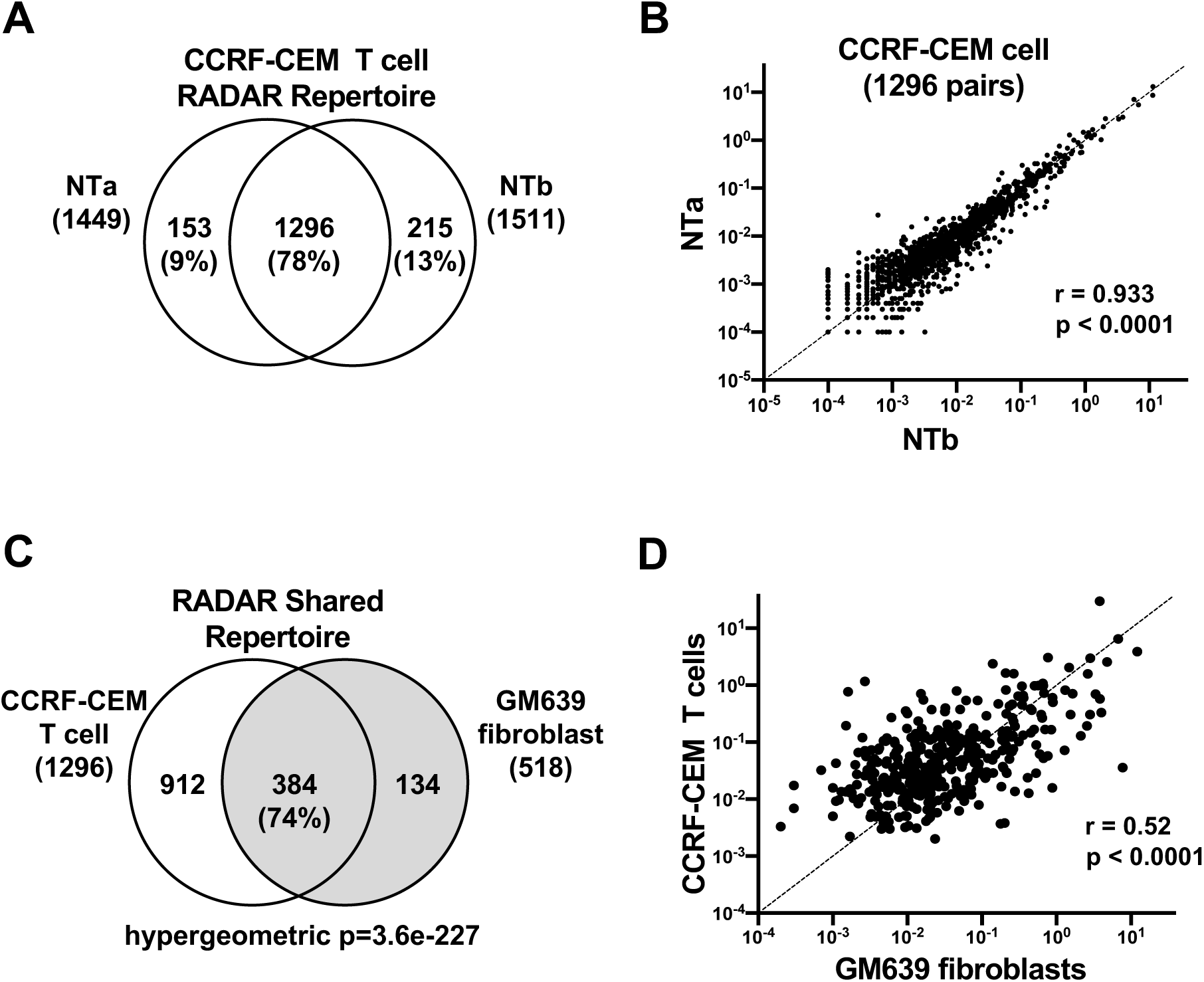
RADAR-MS identifies a limited set of adducted proteins in normally proliferating human cell lines. (A) Venn diagram of shared and unshared proteins in two untreated samples of CCRF-CEM T cells, NTa and NTb, containing 1449 and 1511 proteins, respectively; and a combined repertoire of 1664 non-overlapping proteins. Repertoire size and percent repertoire indicated in parentheses. (B) Pairwise comparison of the IBAQ intensities of the 1296 proteins shared between untreated samples NTa and NTb (Spearman r=0.93; p<0.0001). (C) Venn diagram comparing repertoires of adducted proteins in untreated CCRF-CEM T cells (unshaded) and GM639 fibroblasts (shaded) as determined by RADAR fractionation. Repertoire size and percent of the GM639 repertoire indicated in parentheses. Hypergeometric p=3.6e-227 was calculated based on the following parameters: population size=9000 ([23]; see Methods); sample size A=1296; sample size B=518; set=384; expected successes=75; observed/expected: 175/38; enrichment=5-fold. (D) Pairwise comparison of the intensities of the RADAR shared repertoire of CCRF-CEM T cells and GM639 fibroblasts (Spearman r=0.52; p<0.0001).

We then asked how the repertoire of adducted proteins in CCRF-CEM human T cells compared with the repertoire of adducted proteins in an unrelated cell type, human GM639 fibroblasts. Four untreated samples of GM639 cells were RADAR-fractionated and analyzed by MS, identifying a total of 676 proteins (average 442 per spectrum; range 306-522), with 518 proteins accounting for 99.1% of the MS signal and 77% of the total repertoire (**Table S3**). Comparison of these 518 proteins with the 1296 protein repertoire in CCRF-CEM cells identified 384 shared proteins (**Fig. 2C; Table S3**). These shared proteins accounted for 88% and 85% of the MS signals in the two cell types (**Supplementary Fig. S2D**); and 74% of the GM639 RADAR repertoire, a 5-fold enrichment over 75 common elements expected by chance (hypergeometric p=3.65e-227; **Fig. 2C**). The correspondence of adducts recovered from the two cell types was further evident upon pairwise comparison of their relative abundance (Spearman r=0.52, p<0.0001; **Fig. 2D**). Thus, RADAR-MS identified a limited number of adducted proteins in human cells. We will refer to this repertoire of adducted proteins, based on more than 20 independent RADAR-MS analyses, as the “RADAR shared repertoire”, with the caveat that MS analysis is inherently variable and additional experimentation will be required to compile a truly definitive repertoire.

### 3.4 RADAR enrichment does not correlate with protein abundance

One trivial explanation for repeated identification of a specific protein or class of proteins in independent RADAR-MS analyses is that RADAR fractionation does not effectively eliminate abundant proteins. We therefore asked if the RADAR shared repertoire was enriched in abundant proteins. By intersection of the RADAR shared repertoire (384 proteins) with the CCRF-CEM cell whole cell proteome (6282 proteins, IBAQ>0; http://wzw.tum.de/proteomics/nci60 [22], 342 matching pairs were identified (**Table S4**), which corresponded to 89% of repertoire. Pairwise comparison established that there was no significant correlation between the IBAQ scores of proteins in the RADAR shared repertoire and the whole cell proteome (Spearman r=0.029, p=0.595, **Fig. 3A**).

**Fig. 3.**
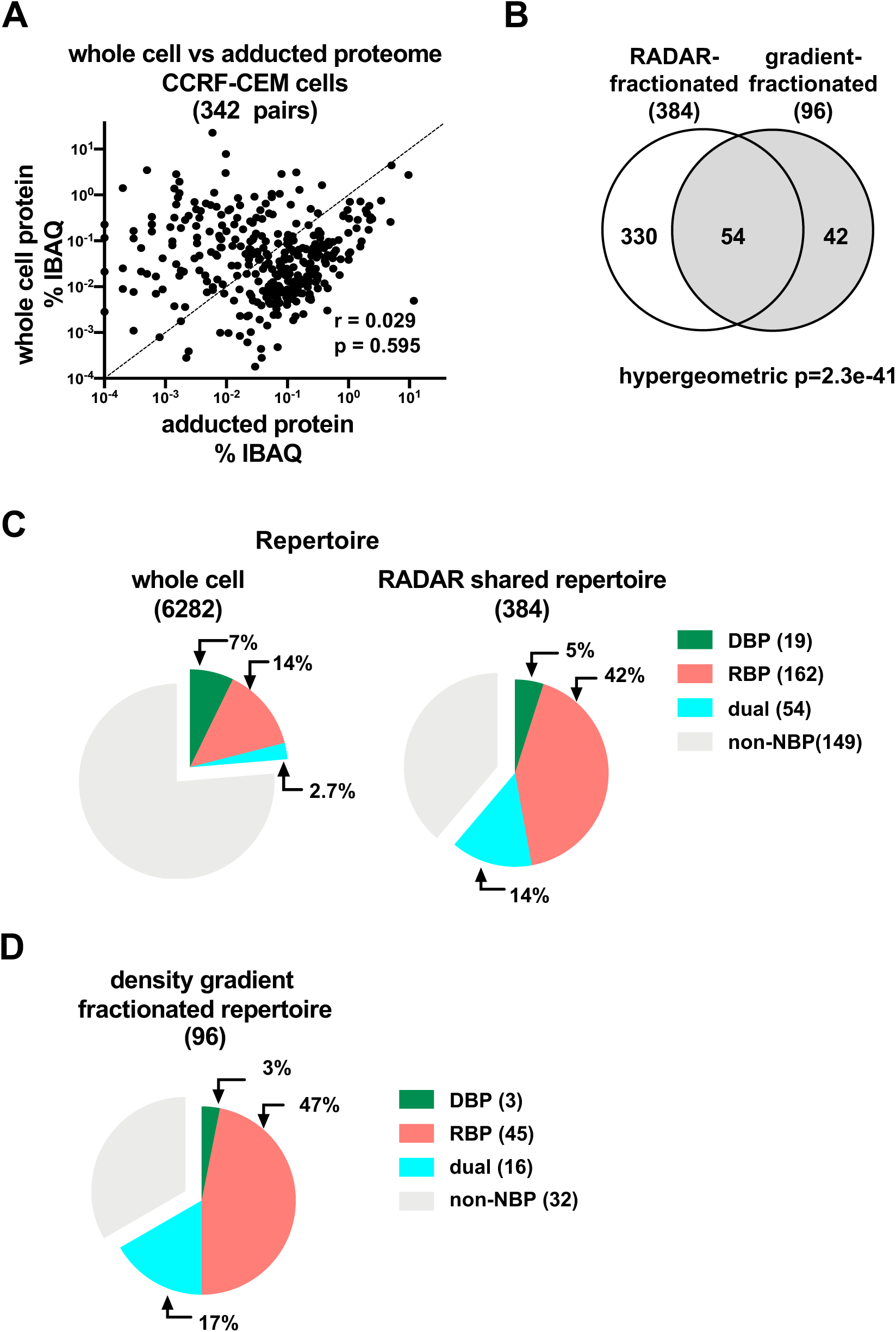
RADAR fractionation enriches RNA binding proteins. (A) Pairwise comparison of the IBAQ intensities of the 342 protein pairs common to the CCRF-CEM whole cell proteome and the RADAR shared repertoire (Spearman r=0.029; p=0.595). (B) Venn diagram illustrating overlap of repertoires of adducted proteins as determined by RADAR fractionation (unshaded) and CsCl buoyant density gradient fractionation (shaded). Repertoire size indicated in parentheses. Hypergeometric p=2.3e-41 was calculated based on the following parameters: population size=6282 sample size A=384; sample size B=96; set=54; expected successes=5.9; observed/expected: 54/5.9; enrichment=9.2-fold. (C) Pie chart indicating the fraction of proteins classified as DBP, RBP or dual binders in the repertoires of the CCRF-CEM whole cell proteome and the RADAR shared repertoire. (D) Pie chart indicating the fraction of proteins classified as DBP, RBP or dual binders in the repertoire determined by buoyant density gradient centrifugation.

### 3.5 Overlap of repertoires of adducted proteins as determined by RADAR and buoyant density gradient fractionation

CsCl buoyant density gradient fractionation has been the standard for isolation of covalent protein-DNA adducts [28]. To compare enrichment of adducts by RADAR fractionation to this classical approach, samples from untreated CCRF-CEM cells which had been lysed in sarkosyl were fractionated by density gradient centrifugation then analyzed by MS, thereby identifying 96 proteins with positive peptide intensity values (**Table S5**). The repertoire of adducted proteins determined by density gradient fractionation exhibited significant overlap with the RADAR shared repertoire (hypergeometric p=2.3e-41, **Fig. 3B**).

### 3.6 RADAR fractionation enriches proteins known to form adducts

Of the 35 human proteins known to form adducts with nucleic acids (Uniprot; **Table S1**), 19 were detected in the whole cell CCRF-CEM proteome [22]. Nine of these proteins (DNMT1, GAPDH, PARP1, PTMA, TOP1, TOP2A, TOP2B, XRCC5 and XRCC6) were identified in the RADAR shared repertoire (**Supplementary Fig. S3A**), where together they accounted for 1.8% of its MS signal (**Table S3)**. This represents a 7.8-fold enrichment over the number of proteins expected in that set by chance (hypergeometric p=5.8e-07). Similarly, the density gradient-fractionated sample (**Table S5**) included five proteins known to form adducts (DNMT1, GAPDH, PTMA, TOP1 and TOP2A), a significant enrichment over the 0.3 proteins expected in that set by chance (hypergeometric p=7.4e-06, **Supplementary Fig. S3B)**. Thus, proteins previously shown to form adducts comprise a small but significant fraction of the RADAR shared repertoire.

### 3.7 RADAR fractionation enriches RNA binding proteins

We further characterized the RADAR shared repertoire by comparing the abundance of nucleic acid binding proteins (NBPs) in it and the publicly available CCRF-CEM whole cell proteome 96282 proteins; [22]). NBP account for 24% of the expressed repertoire of the CCRF-CEM whole cell proteome (**Fig. 3C**), very similar to fraction of the repertoire (20%) that NBP constitute in the reference human proteome (SwissProt, 19,742 entries). RADAR fractionation enriched NBP, which accounted for 61% of the RADAR shared repertoire. Notably, RADAR fractionation especially enriched RNA binding proteins (RBP, 42%) and dual DNA and RNA binding proteins (14%), and modestly depleted DNA binding proteins (DBP, 5%) relative to the whole cell proteome (**Table S6**). Similarly, GO analysis identified as NBP 67% of the proteins in the repertoire of the sample prepared by density gradient fractionation, predominately RBP (47%) and dual binders (17%), along with a small fraction of DBD (3%; **Fig. 3D**).

A considerable number of proteins in the RADAR shared repertoire (149) were classified as non-NBP by GO analysis (**Fig. 3C**). The universe of proteins that bind RNA has undergone recent rapid expansion [29], and the possibility that this might not yet be reflected in GO classifications prompted us to test the possibility that some proteins now known to be RBP might be designated non-NBP in GO classifications. We therefore compared the non-NBP GO species of the RADAR shared repertoire to an experimental proteomics database of RNA-binding proteins recovered from three human cell lines following UV-induced RNA-protein crosslinking (ihRBP, 1753 proteins [7]). Most of the species (79/149; 60%) in the RADAR shared repertoire identified as non-NBP by GO analysis intersected with the ihRBP set (hypergeometric p=1.2e-43; **Supplementary Fig. S3C**). Reclassification of those 79 proteins altered the composition of the RADAR shared repertoire, increasing the fraction of RBP to 63% and reducing the fraction of non-NBP to 18%. This reaffirms the enrichment of RBP among adducted proteins.

### 3.8 Treatment with exogenous formaldehyde increases abundance of adducts present in the repertoires of untreated cells

The results described above support the notion that specific proteins are covalently bound to nucleic acids in normally proliferating human cells, most of which are not known to form adducts as part of their mechanism of action. What might cause these protein-nucleic adducts to form? Formaldehyde is a highly reactive aldehyde that is generated naturally as a product of demethylation reactions in living cells, with concentrations in blood on the order of 0.1 mM [30]. Formaldehyde exhibits clear selectivity in crosslinking [31], making reaction with formaldehyde a plausible source of the limited repertoire of adducted proteins identified.

If endogenous formaldehyde promotes adduct formation by some proteins or classes of proteins, then treatment with exogenous formaldehyde is predicted to increase the abundance of those same proteins or classes of proteins. To test this, we carried out RADAR-MS analysis of CCRF-CEM cells that were treated for 1 hr with 0, 0.5, 1.0 or 2.0 mM formaldehyde. These concentrations are slightly above the endogenous level and in the range of concentrations previously used to study formation and repair of crosslinked proteins in mammalian cells [17, 32, 33], but below levels used to ensure the extensive crosslinking necessary for quantitative immunoprecipitation (5 min, 130 mM formaldehyde [34]). Treatment was carried out in medium lacking or containing 10% fetal bovine serum (Groups A and B, respectively), to provide independent replicates and insights into whether treatment protocols had significant effect on repertoire composition. RADAR fractionation yielded on the order of 265 μg DNA per sample, independent of formaldehyde dose, similar to levels observed in untreated CCRF-CEM samples (**Supplementary Fig. S4A**). The number of proteins detected ranged from 855 to 1081 in the eight samples; and the total MS signal (a sum of intensities of all identified peptides in the sample) was consistent with the number of detected proteins (**Supplementary Fig. S4B; Table S7**). Neither the MS signal nor repertoire size showed an appreciable increase with the formaldehyde dose, indicating that massive chromatin crosslinking did not occur at these formaldehyde concentrations.

To determine which proteins are most reactive with formaldehyde, the proteins that exhibited positive IBAQ values in all treated and untreated samples in both groups were identified, and then the response slopes (the change in relative abundance at each formaldehyde concentration from 0-2 mM) were calculated for each of these 625 proteins using the SLOPE function in Excel (**Table S7**). Response slopes ranged from +1.91 to −2.06. Proteins were ranked based on response slope and the top 100 proteins in each group (16% of total repertoire) compared to identify common proteins (**Fig. 4A**). The two groups included 46 common proteins, far more than the 16 shared elements predicted by chance in the absence of any concentration-dependent increase in adduction in response to treatment with formaldehyde (hypergeometric p=1.8e-15; **Fig. 4A**). Summing up MS signals of these 46 species in each of the two groups established that signals increased in response to increasing formaldehyde, with an average 4-fold increase in both groups at the highest formaldehyde concentration **(Fig. 4B).**

**Fig. 4.**
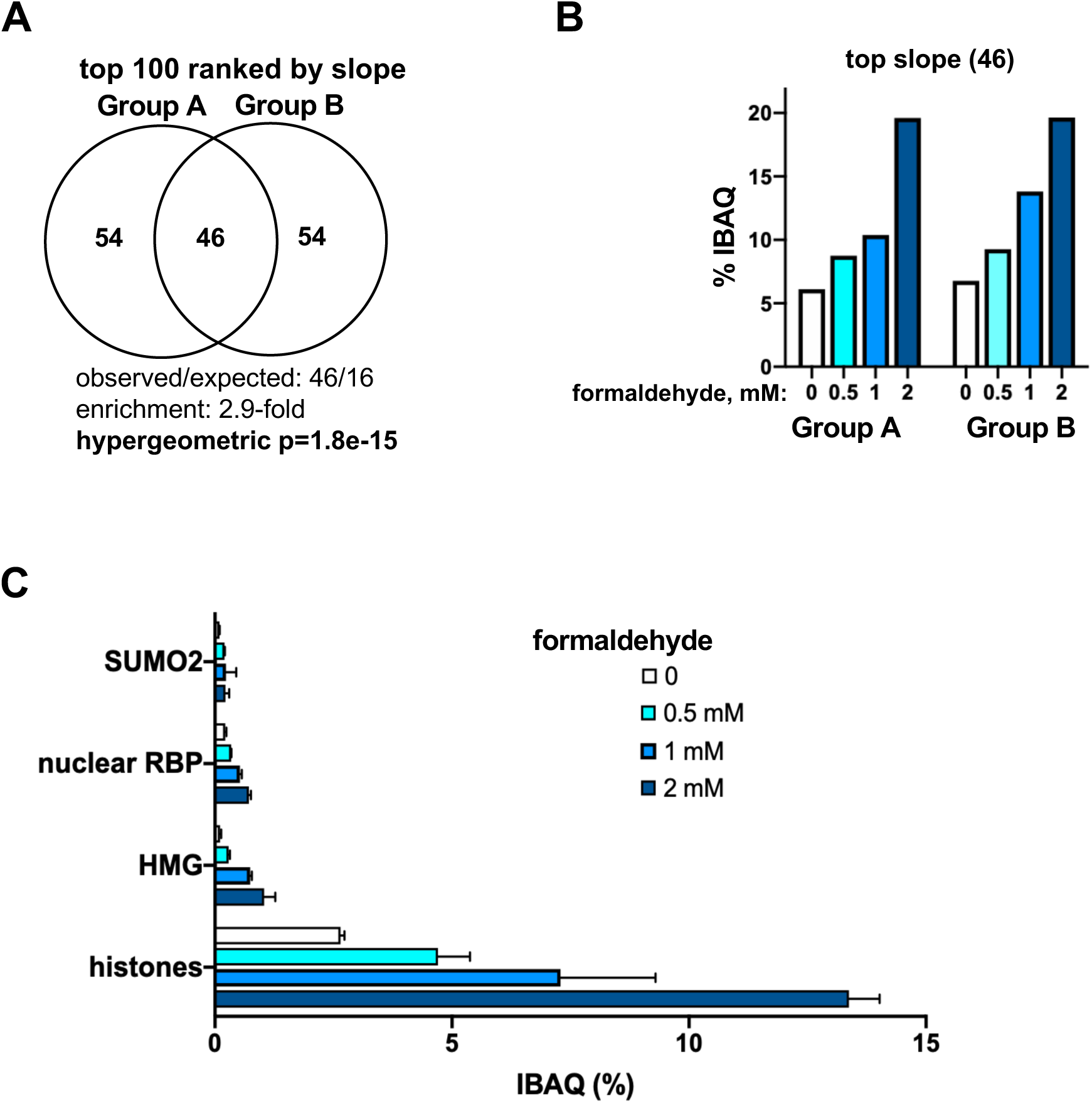
Formaldehyde treatment has little effect on the composition of the repertoire of adducted proteins. (A) Venn diagram of shared and private (unshared) proteins among the 100 proteins with greatest response slopes in Groups A and B. Hypergeometric p=1.8e-15 was calculated based on the following parameters: population size=625 (all proteins with positive intensities); sample size A=100; sample size B=100; set=46; expected successes=16; observed/expected: 46/16; enrichment=2.9-fold. (B) Fractions of MS signal contributed by the 46 shared proteins ranked in the top 100 based upon response slopes. (C) Quantitation of contribution to signal of four proteins/classes of proteins that contributed to 90% of the increased signal at the highest dose of formaldehyde.

The proteins that contributed to 90% of the increased signal at the highest dose of formaldehyde belonged to four classes: histones, HMG proteins, nuclear RBP, and SUMO2 **(Fig. 4C**). The increased signal of SUMO2 may reflect posttranslational modification that targets proteins for proteolytic repair, as was also seen in RADAR-SILAC analysis of etoposide-treated cells (**Fig. 1C**). Thus, a limited repertoire of proteins appears to be highly reactive to exogeneous formaldehyde at doses only slightly surpassing its physiological level.

### 3.9 The RADAR shared repertoire is enriched for proteins with specific functions

The distribution of proteins of the RADAR shared set by relative abundance showed that a very limited number of species accounted for most of the MS signal in each of the two cell types **(Fig. 5A)**: more than 50% of the signal could be assigned to a total of 13 proteins, and more than 75% of the signal to 50 proteins. Based on GO analysis, RBP accounted for most of the MS signal in the RADAR shared repertoire (65%; **Fig. 5B**). This represents at least a 2.2-fold enrichment over the whole cell proteome, a minimum estimate because not all RBPs are scored by GO classification (e.g. **Supplementary Fig. S3C**).

**Fig. 5.**
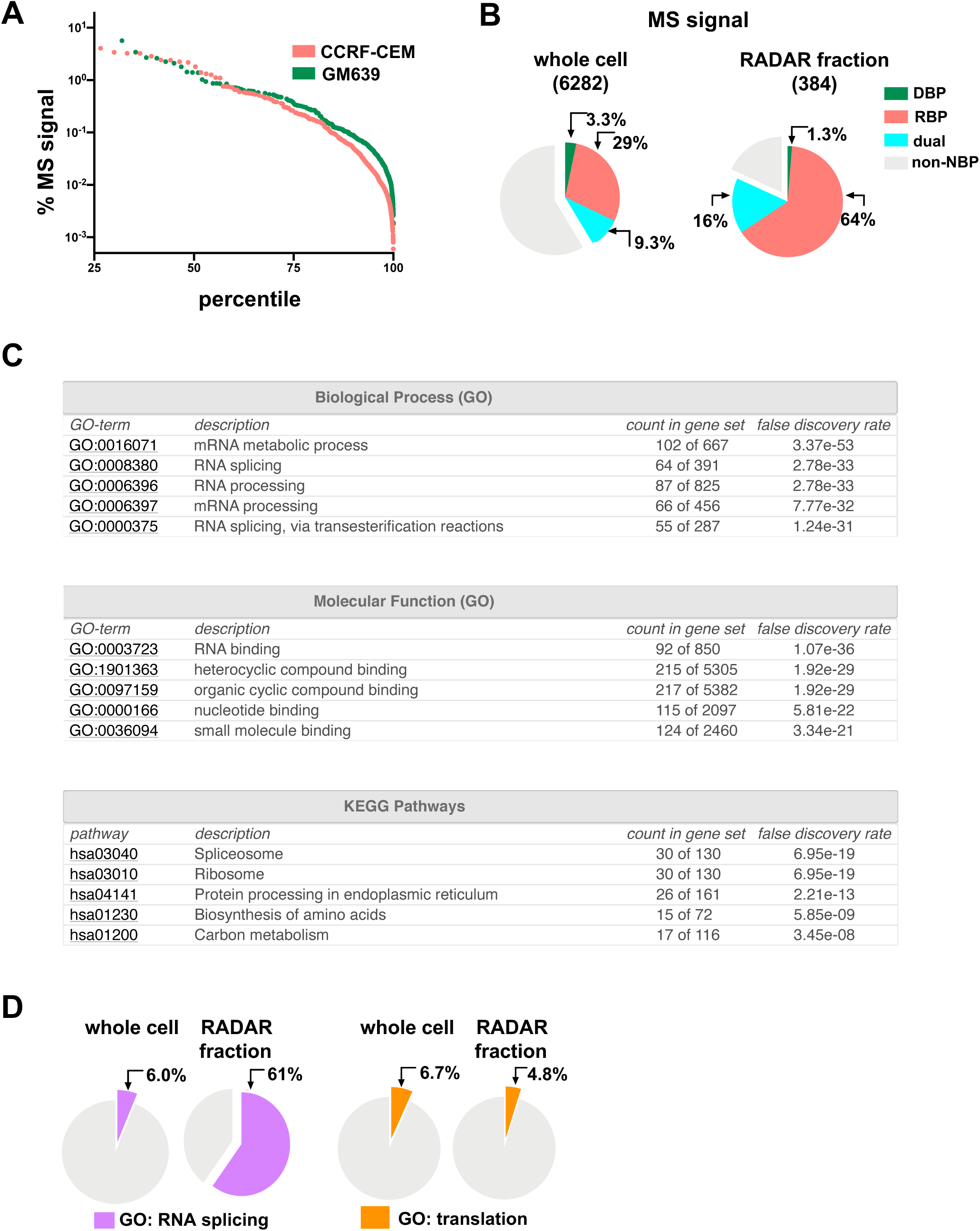
Proteins enriched in the repertoire of adducted proteins. (A) Frequency distribution of IBAQ scores in the RADAR shared repertoire of CCRF-CEM T cells and GM639 fibroblasts. (B) Pie chart indicating the fraction of MS signal contributed by proteins classified as DBP, RBP or dual binders in the CCRF-CEM whole cell proteome and the RADAR shared repertoire. (A pie chart based on repertoire rather than MS signal is presented in **Fig. 3C**.) (C) Highly significant functional enrichments in the RADAR shared repertoire based on analysis of protein-protein interaction networks (STRING database). (D) Pie chart indicating the fraction of MS signal contributed by proteins associated with the GO terms RNA splicing and translation in the CCRF-CEM whole cell proteome and the RADAR shared repertoire.

Analysis of protein-protein interaction networks identified specific GO terms for biological processes associated with the adducted proteome with very high significance (FDR<e-30): mRNA metabolism, RNA splicing, RNA processing, mRNA processing and RNA splicing via transesterification reactions (**Fig. 5C**). Molecular function GO terms (FDR<e-21) and KEGG pathways (FDR<e-08) provided further support for associations of these proteins with RNA processing and biogenesis. Strikingly, proteins characterized by a single GO term, RNA splicing, comprised 61% of the MS signal of adducted proteins, a 10-fold increase over the whole cell proteome (**Fig. 5D**, left). In contrast, MS signals of proteins characterized by the GO term translation were not enriched among adducted proteins (**Fig. 5D**, right). Thus the strong associations with splicing appear to reflect specific, nuclear location-related, rather than general, RNA-related activities of abundant proteins in the RADAR fraction.

## Discussion

The results described here demonstrate that proteins adducted to nucleic acids in human cells can be discovered and identified by combining RADAR fractionation with MS. Application of this approach to normally proliferating human cells identified a shared repertoire of adducted proteins with a quantitative profile that was significantly concordant in two unrelated cell types, CCRF-CEM T cells and GM639 fibroblasts (Spearman r=0.52; p<0.0001; **Fig. 2D**). The repertoire determined by RADAR fractionation corresponded very well with that determined by the classical approach of buoyant density gradient ultracentrifugation (hypergeometric p=2.3e-41; **Fig. 3B**). Because density gradient centrifugation and RADAR use distinct approaches to separate nucleic acids from non-covalently bound proteins, the subset of adducted proteins identified by both methods is likely to contain bona fide covalent nucleoprotein complexes and not fractionation artifacts.

MS analysis is inherently variable and the repertoire of adducted proteins as determined thus far was not intended to be definitive, even though it was based on more than 20 independent RADAR-MS analyses. Nonetheless, its composition provides experimental insights into the classes of proteins that form adducts in normally proliferating cells.

The repertoire of adducted proteins — or the “adductome” — contained a relatively limited number of proteins. A minor but significant fraction of the adducted repertoire (hypergeometric p=5.8e-07, 1.8% MS signal) was comprised of proteins for which formation of adducts with DNA is key to mechanism of action, among them TOP1, TOP2, DNMT1 and Ku (XRCC5/6). Histones and HMG proteins were considerably enriched among adducted proteins, along with nuclear RNA binding proteins, especially proteins associated with mRNA splicing (**Fig. 5C**). Abundant RNA binding proteins with other functions were not enriched, including proteins associated with translation. Adduction of proteins involved in mRNA splicing may reflect primary sequence, as only some amino acid residues can form adducts. It is also an intriguing possibility that RNA splicing proteins form crosslinks with their substrates to facilitate the long-range interactions necessary for RNA processing.

Enrichment of RBP in the adductome was somewhat unexpected, as the majority of adduct-forming proteins described thus far form crosslinks with DNA. This may reflect more intensive study of protein-DNA adducts, which may be more cytotoxic than protein-RNA adducts; or it may be a consequence of methods used to fractionate adducted proteins. The buoyant density of RNA is slightly greater than that of DNA, and a density gradient protocol designed to enrich protein-DNA adducts may not efficiently recover protein-RNA adducts (e.g. [35]), while these will be recovered by RADAR fractionation.

Treatment of cells with low concentrations of formaldehyde increased the abundance in the adductome of histones, HMG proteins and nuclear RBD, the same classes of proteins enriched in normally proliferating cells. This is consistent with the possibility that endogenous formaldehyde generated by ongoing metabolism promotes adduct formation in normally proliferating cells. Formaldehyde is a highly reactive aldehyde that is generated naturally as a product of demethylation reactions in living cells, reaching concentrations in blood in the range of 0.1 mM even in the absence of environmental exposure [30]. Formaldehyde exhibits clear selectivity, crosslinking histones but failing to crosslink other proteins, including serum albumin and even the formidable DNA binding protein, lactose repressor [31]. This selectivity in protein crosslinking makes formaldehyde an especially plausible candidate as the agent that promotes formation of a limited repertoire of protein-nucleic acid adducts in normally proliferating cells. Alcohol dehydrogenase 5 (ADH5) normally eliminates formaldehyde in mammalian cells, but upon impairment of this pathway endogenous formaldehyde can become a genotoxin and carcinogen [36]. This highlights the importance of adduct repair in maintaining genomic structure and provides a further alert to the contribution of repair insufficiencies to cancer.

RADAR-MS analysis may be useful for understanding the pathways that repair potentially cytotoxic adducts. RADAR fractionation is rapid, high throughput and cost-effective, and requires only standard laboratory equipment. RADAR lysis in chaotropic salts interrupts proteolytic processing of repair intermediates, so the peptides identified by MS provide a snapshot of ongoing repair. This was evident as enrichment of peptides surrounding the active site tyrosine in the analysis of TOP1 (**Fig. 1D**), probable intermediates of proteolytic repair of the covalently bound polypeptide. Post-translational sumoylation frequently marks a protein for proteolytic degradation, and the observed enrichment of SUMO2 in etoposide-treated CCRF-CEM cells (**Fig. 1C**) and in extracts of formaldehyde-treated cells (**Fig. 4C**) provides further evidence of the ability of RADAR-MS to detect such intermediates. Proteolytic repair intermediates may evade immunodetection because they lack epitopes critical for antibody recognition. Moreover, the RADAR-MS approach we have developed for adduct characterization can be readily applied in a variety of contexts to identify repair intermediates and provide information on how key factors participate in adduct repair.

## Author contributions

K.K. and N.M. designed the experiments. K.K. carried out the experiments. KK and NM wrote the manuscript.

## Acknowledgments

We thank Shao-En Ong and Emily Myers for invaluable advice and assistance with MS. This research was supported by NIH P01 CA077852 (to N.M.) and NIH R21 CA194876 (to N.M. and S.E. Ong). This research used an EASY-nLC1200 UHPLC and Thermo Scientific Orbitrap Fusion Lumos Tribrid mass spectrometer purchased with funding from a National Institutes of Health SIG grant S10OD021502 to S.E. Ong.

## Appendix A. Supplementary data

Supplementary material related to this article can be found, in the online version.

## Supplementary Figure Legends

**Fig. S1.**
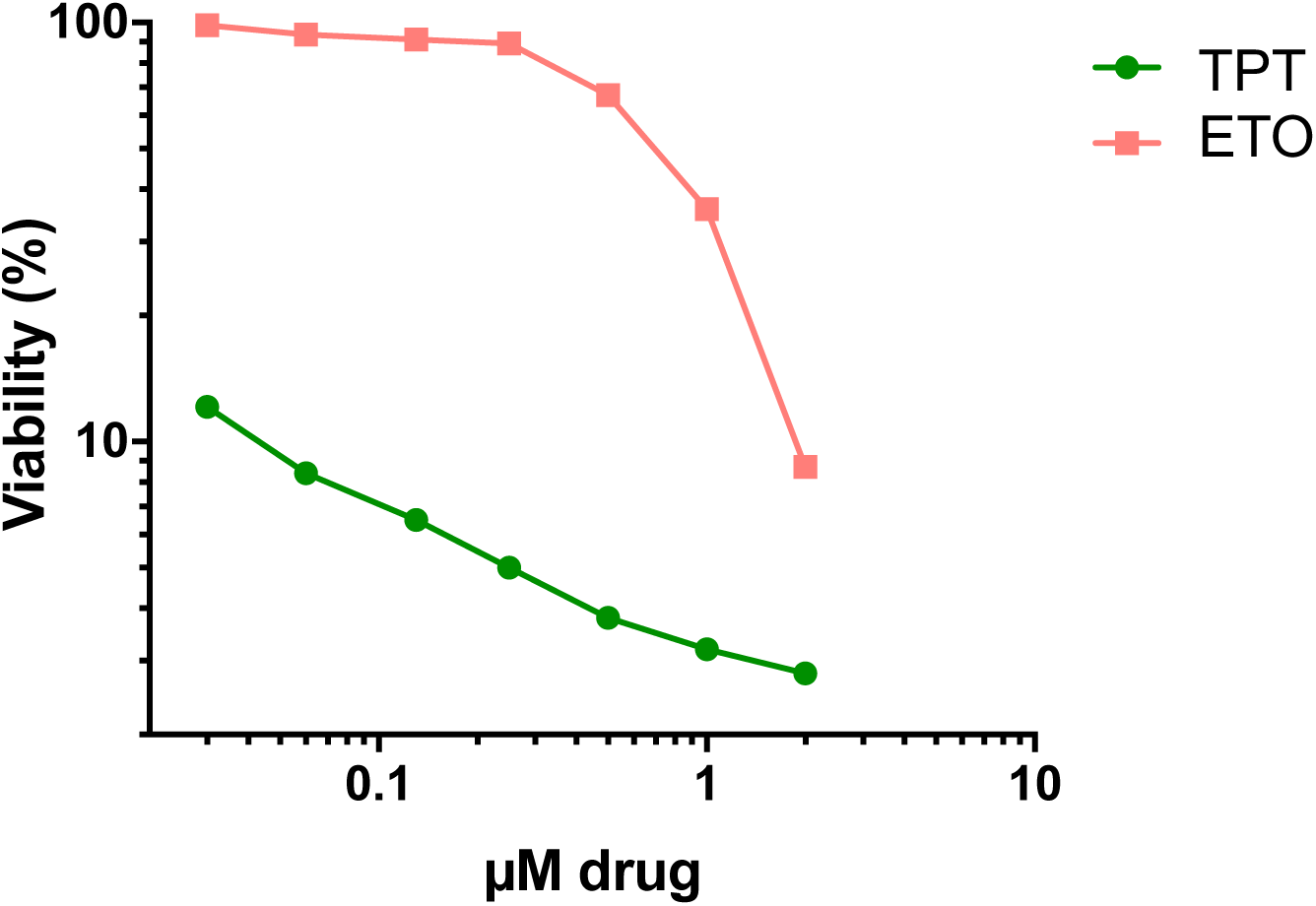
Sensitivity of CCRF-CEM cells to topoisomerase inhibitors. CCRF-CEM cells were treated with indicated concentrations of topotecan (TPT) or etoposide (ETO), which poison TOP1 and TOP2, respectively. The viability was assayed by the CellTiter-Glo^®^ ATPase assay (Promega) after 72 hours incubation with the drug.

**Fig. S2.**
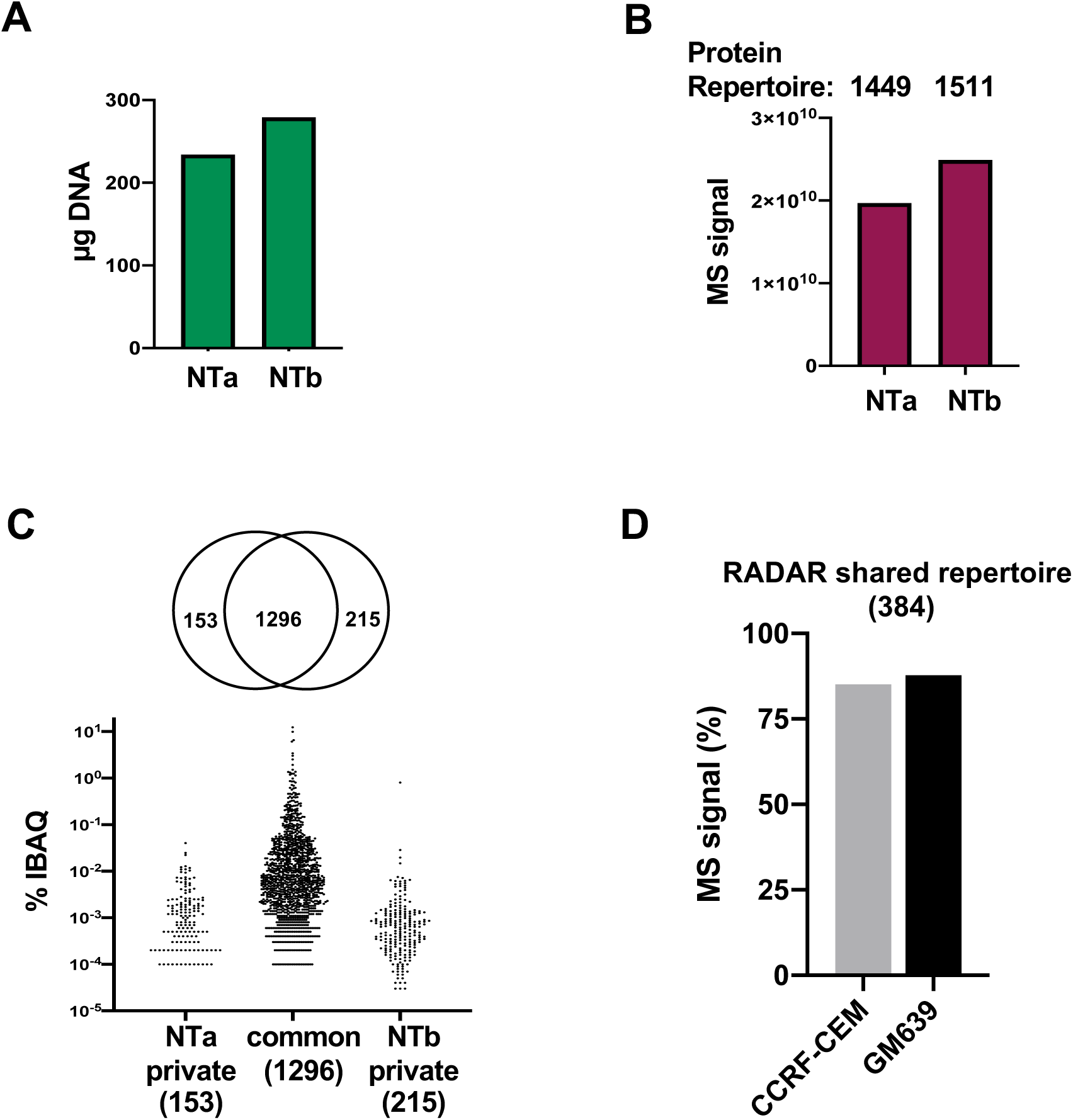
Recovery of adducted proteins after RADAR fractionation. (A) DNA content of RADAR-fractionated CCRF-CEM cells from untreated samples NTa and NTb. (B) Total IBAQ intensity of peptides (y-axis) and size of protein repertoires in samples NTa and NTb. (C) *Above*, Venn diagram showing overlap of proteins common to samples NTa and NTb or found only in one of those two samples (private). *Below*, percent contribution to IBAQ of proteins in each of these groups. Numbers of proteins in each category shown in parentheses. (D) Percent MS signal contributed to the RADAR shared repertoire (by adducted proteins identified in CCRF-CEM or GM639 cells.

**Fig. S3.**
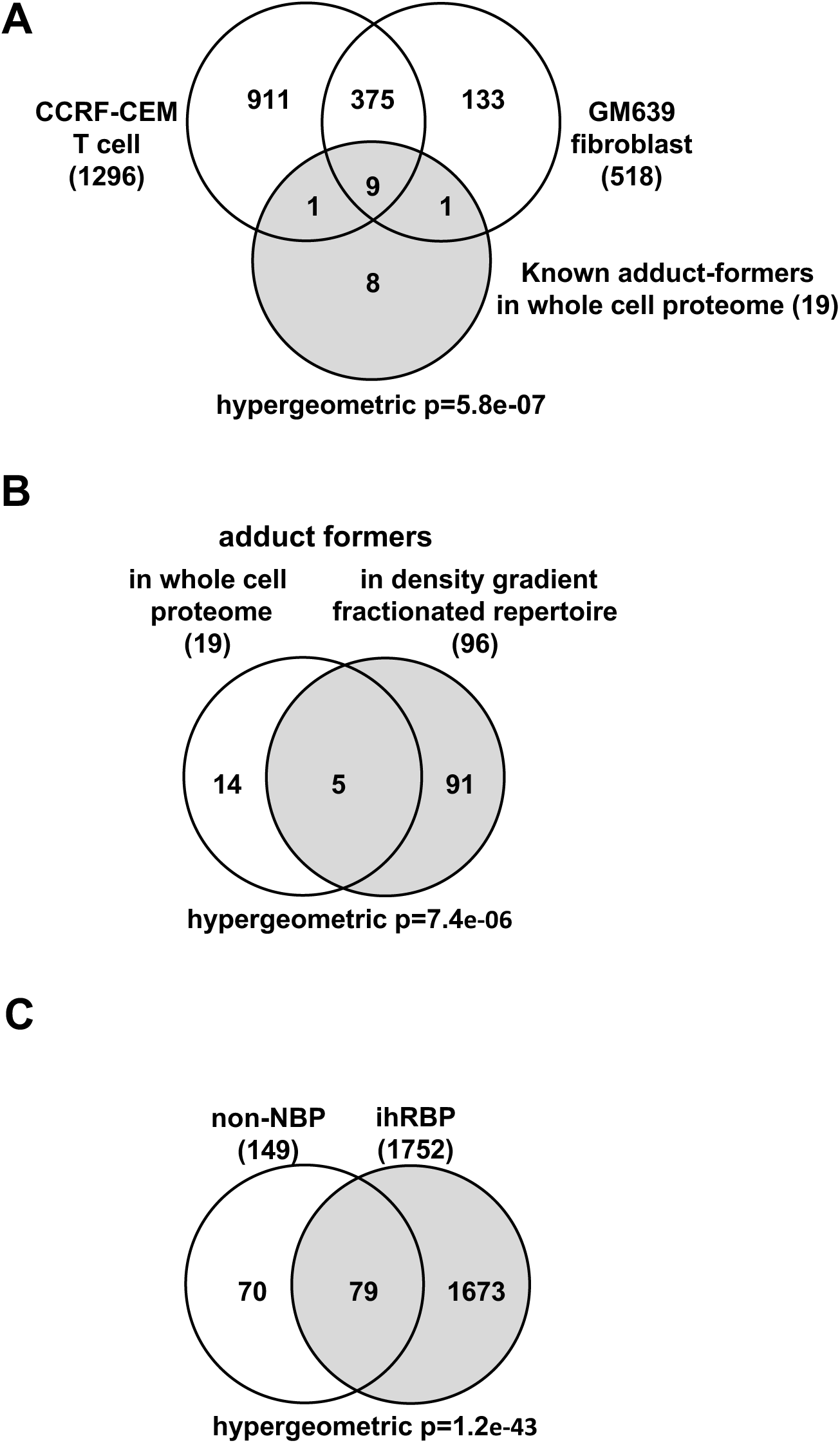
Recovery of adducted proteins after RADAR fractionation. (A) Venn diagram of overlaps between known adduct forming proteins in the CCRF-CEM T cell whole cell proteome (gray shading) and the RADAR repertoires of CCRF-CEM and GM639 cells (unshaded). Repertoire sizes indicated in parentheses. Hypergeometric p=5.8e-07 for the intersect of the RADAR shared repertoire (384) and known adduct formers (19) was calculated based on the following parameters: population size=9000 ([23]; see Methods); sample size A=384; sample size B=19; set=9; expected successes=1.2; observed/expected: 9/1.2; enrichment=7.7-fold. (B) Venn diagram of overlap between known adduct forming proteins in CCRF-CEM T cell whole cell proteome and the repertoire of adducted proteins as determined by density gradient fractionation followed by MS (gray shading). Repertoire size indicated in parentheses. Hypergeometric p=7.4e-06 was calculated based on the following parameters: population size=6282; sample size A=19; sample size B=96; set=5; expected successes=0.29; observed/expected: 5/0.29; enrichment 17-fold. (C) Venn diagram of overlap between non-NBP identified by GO analysis of the RADAR shared repertoire and the ihRB proteome ([7]; gray shading). Repertoire size indicated in parentheses. Hypergeometric p=1.2e-43 was calculated based on the following parameters: population size=19742 (Uniprot); sample size A=149; sample size B=1752; set=79; expected successes=13.2; observed/expected: 79/13.2; enrichment=6.0-fold.

**Fig. S4.**
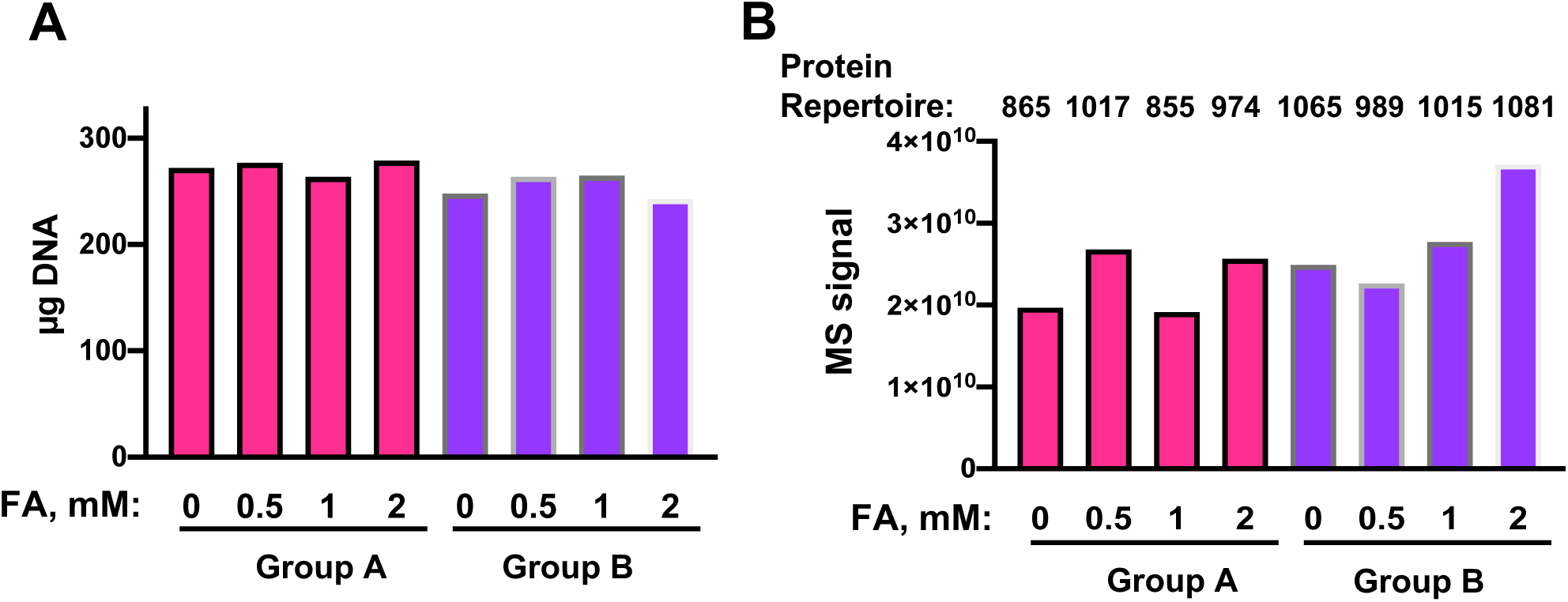
Repertoires of adducted proteins in CCRF-CEM T cells treated with formaldehyde. (A) DNA content of RADAR-fractionated CCRF-CEM cells from Groups A and B, treated with 0, 0.5, 1.0 or 2.0 mM formaldehyde (FA) in the absence or presence of serum, respectively. (B) Total IBAQ intensity of peptides (y-axis) and size of protein repertoires in samples described in panel A.

## Supplementary Tables

**Table S1**. 35 known adduct forming proteins.

**Table S2**. RADAR-MS analysis of CCRF-CEM cells.

**Table S3**. CCRF-CRM and GM639 shared repertoire.

**Table S4**. RADAR shared repertoire vs CCRF-CEM whole cell repertoire.

**Table S5**. Density gradient-MS analysis of adducts in CCRF-CEM cells.

**Table S6**. NBP GO enrichments.

**Table S7**. Formaldehyde response of CCRF-CEM cells.

